# Complementary inhibitory weight profiles emerge from plasticity and allow attentional switching of receptive fields

**DOI:** 10.1101/729988

**Authors:** Everton J. Agnes, Andrea I. Luppi, Tim P. Vogels

**Affiliations:** Centre for Neural Circuits and Behaviour, University of Oxford, OX1 3SR Oxford, United Kingdom

**Author notes:** Department of Clinical Neurosciences, University of Cambridge, Cambridge, UK.

## Abstract

Cortical areas comprise multiple types of inhibitory interneurons with stereotypical connectivity motifs, but their combined effect on postsynaptic dynamics has been largely unexplored. Here, we analyse the response of a single postsynaptic model neuron receiving tuned excitatory connections alongside inhibition from two plastic populations. Depending on the inhibitory plasticity rule, synapses remain unspecific (flat), become anti-correlated to, or mirror excitatory synapses. Crucially, the neuron’s receptive field, i.e., its response to presynaptic stimuli, depends on the modulatory state of inhibition. When both inhibitory populations are active, inhibition balances excitation, resulting in uncorrelated postsynaptic responses regardless of the inhibitory tuning profiles. Modulating the activity of a given inhibitory population produces strong correlations to either preferred or non-preferred inputs, in line with recent experimental findings showing dramatic context-dependent changes of neurons’ receptive fields. We thus confirm that a neuron’s receptive field doesn’t follow directly from the weight profiles of its presynaptic afferents.

---

Inhibitory neurons exhibit large variability in morphology, connectivity motifs, and electrophysiological properties^1–4^. Inhibition often balances excitatory inputs, thus stabilising neuronal network activity^5,6^ and allowing for a range of different functions^7–11^. When both inhibitory and excitatory inputs share the same statistics and their weight profiles are similar^12^, the resulting state of the post-synaptic neuron is one of precise balance of input currents^9^. Modulation of inhibition, e.g., a decrease or increase in local inhibitory activity and, consequently, a change in the balance between excitation and inhibition, can control the activity of neuronal groups^13,14^, and it is believed that disinhibition is an important mechanism for the implementation of high-level brain functions, such as attention^14,15^, memory retrieval^6,16,17^, signal gating^18,19^, and rapid learning^11^. The state of balance is thought to be achieved and maintained by inhibitory plasticity, e.g., a Hebbian-like inhibitory plasticity rule^6^ (increase in synaptic weights for correlated pre- and postsynaptic activity), as observed in auditory cortex^20^. Other types of inhibitory plasticity have also been observed, such as a form of anti-Hebbian inhibitory plasticity (decrease of synaptic weights for correlated pre- and postsynaptic activity)^21–23^ that has been proposed as a mechanism for memory formation^24^.

Given that cortical circuit motifs feature multiple interneuron types^1,2,7^, we wondered how these opposing types of plasticity may act in concert on the same postsynaptic target, and how the resulting synaptic weight profiles can help to shape the receptive field of the postsynaptic neuron. We speculated that two plasticity rules could form complementary synaptic weight profiles for inhibitory connections, such that synapses following a Hebbian-like inhibitory plasticity rule would mirror excitatory inputs; Anti-Hebbian plasticity should impose strong inhibitory inputs for weak excitatory ones, and vice-versa. Such opposite wiring profiles of distinct inhibitory synapse populations are in line with intracellular recordings showing that strong inhibitory postsynaptic potentials can be elicited by stimuli with preferred orientations of the postsynaptic neuron^25,26^, but also by stimuli with non-preferred orientations^27,28^. What’s more, dynamically changing receptive fields could be achieved through targeted modulation of a specific type of inhibition.

Altered receptive properties have been widely observed, e.g., in mouse auditory cortex where neurons change their preferred sound frequency with varying sound intensity^29^. In macaque primary visual cortex (V1) neurons can modulate their response according to an extra cue of a different (auditory) sensory nature^30^. Intriguingly, they responded either more strongly to their preferred stimulus or, on the contrary, they were more suppressed when a pure tone was played alongside the presentation of the visual stimuli. In macaque V4^31^ and V5^32^ neurons have been shown to change how they represent different stimuli during detection and discrimination tasks, and in macaque V4^33^ some neurons change their hue preference when subjected to single-hue or naturally coloured images. Finally, recent work by Billeh *et al*. ^34^ showed that in mice, visual neurons change their response to the direction of motion of visual stimuli depending on either the temporal of the spatial frequency of the stimulus (drifting grating). These results suggest that the receptive fields of sensory neurons are dramatically affected by input, context or attentional state, but it is unclear by which mechanisms such changes can transpire.

Here, we tested how the response of a single neuron is affected when the activity of presynaptic inhibitory populations is modulated. We combined two hypotheses to address this question. First, we considered that different types of inhibitory interneurons may follow distinct synaptic plasticity learning rules (Fig. 1A), thus creating different connectivity profiles onto postsynaptic neurons^9^ (Fig. 1B) such as those observed for e.g., parvalbumin-positive (PV+) and somatostatin-positive (SOM+) interneurons^35^. PV+ interneurons may follow a Hebbian-like plasticity rule^6,20^, thus targeting pyramidal neurons with similar preferred orientation^35^. SOM+ interneurons, on the other hand, could follow a non-Hebbian plasticity rule (e.g., anti-Hebbian or homeostatic), which results in a non-selective connectivity^35^. Our second hypothesis posits that changes in the activity of inhibitory neurons are responsible for the highly variable receptive fields observed in recent experiments^29–34^ (Fig. 1C). This hypothesis extrapolates from evidence of cortical disinhibition during functional tasks^13,36^, and requires that a different brain region provide attentional or contextual signals, such as observed in prefrontal cortex and regions in the frontal lobe^37–40^.

**FIG. 1.**
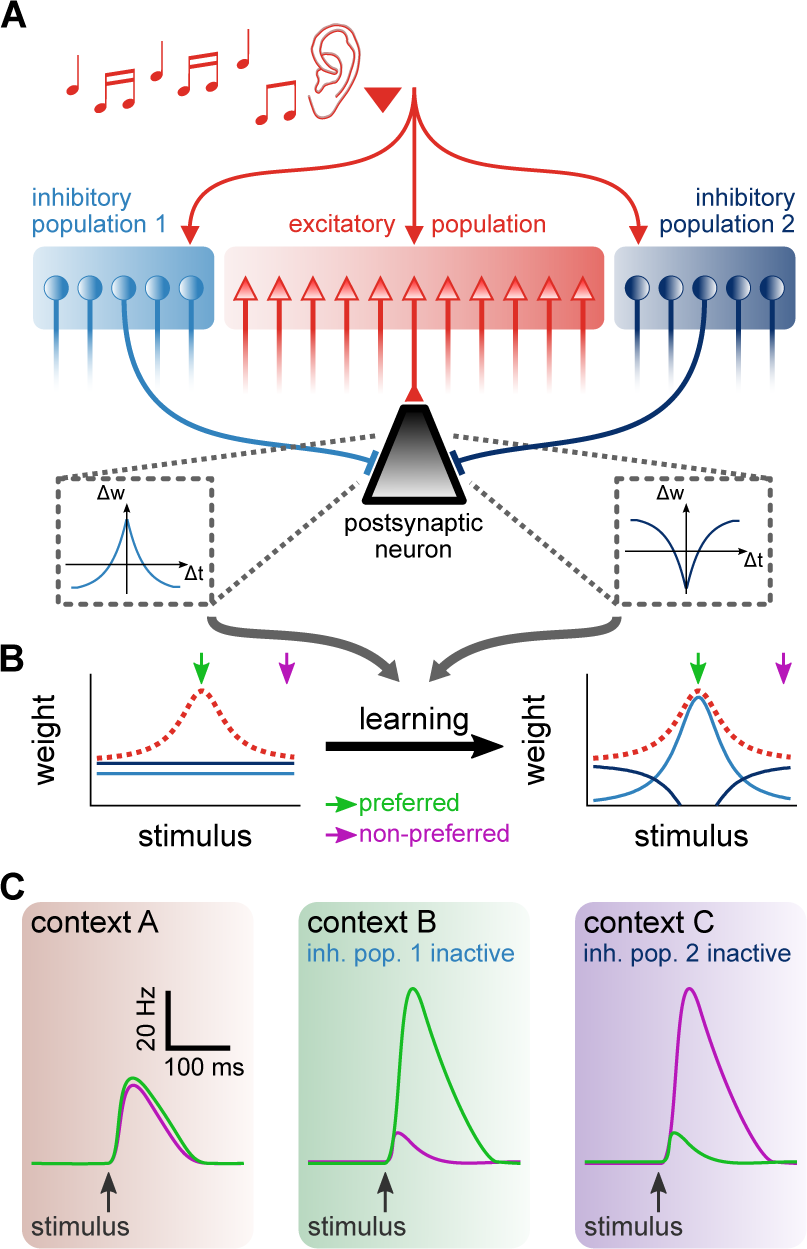
Learning of two distinct inhibitory populations and postsynaptic response due to attentional switch between contexts. **A**, Schematic of co-active plasticity rules. A postsynaptic neuron (black triangle) receives tuned excitatory input (red population) and inhibition from two distinct populations (blue populations). The two inhibitory populations follow different synaptic plasticity rules. ∆*w* indicates change in synaptic weight and ∆*t* indicates interval between pre- and post-synaptic spikes. **B**, Initially un-tuned inhibitory weights (blue lines) acquire different synaptic weight profiles after learning that depend on the excitatory weight profile (red dashed line). **C**, Contextual changes (e.g., due to attention), which we hypothesise to be responsible for modulating the activity of inhibitory populations result in different postsynaptic responses to the same stimulus^29–34^, such that preferred (green) and non-preferred (purple) stimuli elicit postsynaptic responses with different amplitudes.

To investigate the origins of such varying responses from the same cell, we investigated the behaviour of a single postsynaptic neuron model receiving tuned excitatory inputs, and inhibition from two distinct populations. Input tuning may correspond to preference to a specific sound frequency^12^, orientation of visual cues^41^ or to colour hue^33^, taste^42^, whisker stimulation^43^, or position in space^44^. We show that when distinct biologically plausible plasticity rules operate on the synapses of different inhibitory populations, at least three different tuning profiles may emerge. After learning, the postsynaptic neuron arrives at a balanced state with respect to its excitatory and inhibitory inputs. In this state, preferred signals are transiently revealed, but steady state responses are indiscriminate of the stimulus preference^6^ (i.e., its ‘orientation’, etc.), regardless of the inhibitory connectivity. However, we could substantially alter the responses of the postsynaptic neuron by modulating the activity of either of the two presynaptic inhibitory populations, allowing for the propagation of facets of the input patterns that were previously quenched by inhibition. Such inhibitory modulation can thus serve as a mechanism to selectively filter stimuli according to, e.g., attentional cues, as observed in recent experiments^29–34^. In summary, our work proposes a simple biological implementation for an attentional switch of input selectivity, and provides a solution for how such a neuronal circuit can emerge with autonomous and unsupervised, biologically plausible plasticity rules. To our best knowledge, our model is the first proof of principle that the receptive field of a neuron, i.e., its response to presynaptic stimuli, must not follow directly from the (excitatory) presynaptic weight profiles.

## Results

To study the effect of interacting populations of feedforward inhibition, we investigated the response of a single postsynaptic leaky integrate-and-fire neuron receiving tuned excitatory inputs and inhibition from two distinct populations. Excitatory inputs were organised into a single population, subdivided into 16 signal groups of 200 excitatory afferents. Inhibitory inputs initially formed a single population, mirroring the excitatory subdivision, but with 50 afferents per group. Subsequently, we split the inhibitory inputs into two populations with 25 afferents per signal group (Fig. 2A, see Methods for details), allowing us to obtain two differently tuned populations (presumably *types*) of inhibition. Excitatory and inhibitory afferents belonging to the same group shared temporal fluctuations in firing rates, termed input patterns, even if they belonged to different populations. In our simulations, input patterns could either be *natural* or *pulse*. Natural inputs were generated through an inhomogeneous Poisson process based on a modified Ornstein-Uhlenbeck process (Fig. 2B,C), such that neurons of the same signal group also had temporally-correlated firing patterns (Fig. 2C, top; see also Ujfalussy *et al.* ^45^). The resulting long-tail distribution of inter-spike-intervals (Fig. 2C, bottom) was similar to experimentally observed spike patterns *in vivo*^46,47^. We used this type of input to train inhibitory synapses via plasticity rules, and to quantify steady-state (average) postsynaptic responses.

**FIG. 2.**
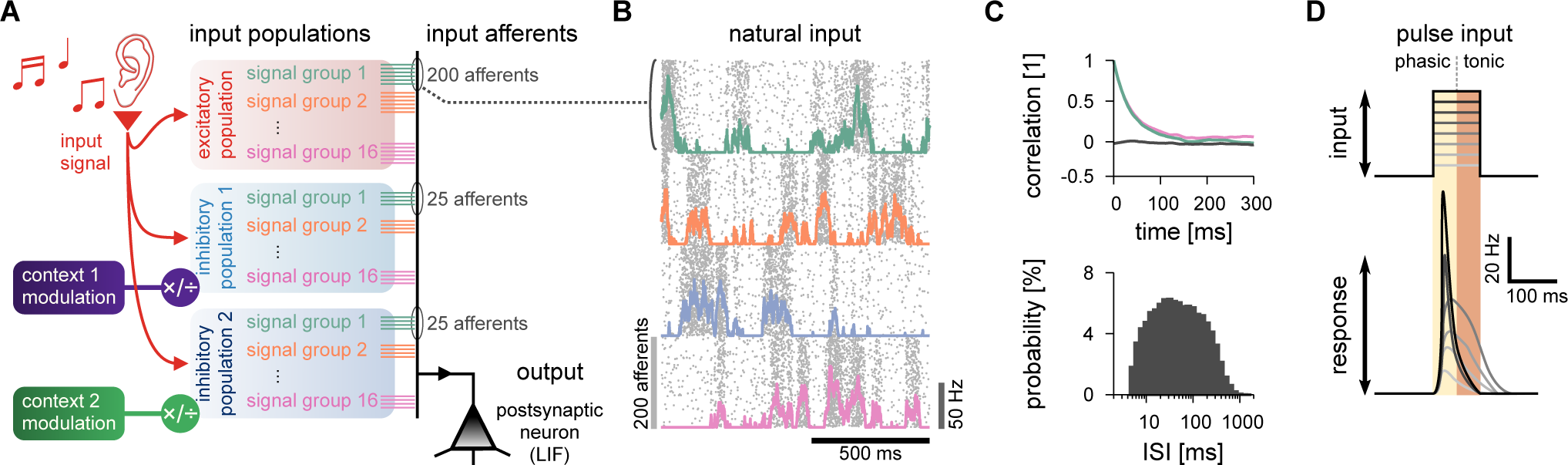
Model details. **A**, Schematic of the input organisation. An external signal (representing, e.g., sound) was delivered through three input populations (one excitatory and two inhibitory), with 16 input signals per population (representing, e.g., sound frequency). Each signal was simulated by 250 independent, but temporally correlated, spike trains (input afferents); 200 excitatory, and 50 inhibitory divided into two groups of 25. One postsynaptic neuron (black triangle) was the output of this system, simulated as a single-compartment leaky integrate-and-fire neuron (LIF). The firing rate of each of the inhibitory populations was modulated by a contextual cue (green and purple boxes). Excitatory and inhibitory input spike trains were generated as point processes (see Methods for details). **B**, Natural input statistics. Raster plot (grey dots) of 800 neurons that take part in 4 signal groups (200 neurons per signal group), each with firing-rate changing according to a modified Ornstein-Uhlenbeck process (coloured lines; Methods). **C**, Temporal autocorrelation (top) and distribution of the inter-spike intervals (ISI; bottom) of the pre-synaptic inputs. The autocorrelation of two groups are shown (green and pink), as well as the correlation between two different groups (black). Autocorrelation is computed as the Pearson coefficient with a delay (x-axis; Methods). **D**, Pulse input schematic. A step-like increase in the firing rate of a given input group lasting 100 ms with varying firing rates (grey scale). The postsynaptic response is separated in *phasic* (first 50 ms), and *tonic* (last 50 ms).

In the alternative pulse input regime we analysed transient responses with 100-ms long pulses of varying amplitudes^6^. Pulses were delivered through a single signal group of excitatory and inhibitory afferents, while all other groups remained at baseline firing-rate (Methods). Responses were quantified according to postsynaptic firing rates during the first (phasic) and last (tonic) 50 ms stimulation (Fig. 2D), averaged over 100 trials. Separating responses in phasic and tonic allowed us to discriminate changes in output due to the input onset, and slower integration of the pulse, respectively.

Learning was implemented via three distinct inhibitory plasticity rules (Fig. 3), in three different combinations. We first implemented a Hebbian rule (that potentiated synaptic weights for coincident pre- and postsynaptic spikes and depressed them for sole presynaptic spikes^6^; Fig. 3A) in one of the two inhibitory populations, while the synapses of the other inhibitory and the excitatory population remained fixed. This learning rule has previously been shown to generate inhibitory weight profiles that mirror the excitatory synaptic weight profiles of a postsynaptic neuron, imposing a firing-rate fixed-point (target; Fig. 3A) by balancing excitation and inhibition^6^, supporting similar experimental findings in mouse auditory cortex^20^. Next, we implemented the Hebbian plasticity rule in one of them and a *scaling* plasticity rule (Fig. 3B) in the other population. The homeostatic scaling rule up- or down-regulates the entire synapse population to reach a predetermined target firing-rate. Notably, this plasticity rule was purely local, taking only synaptic weights and postsynaptic firing rate into account, similarly to the experimentally observed scaling of inhibitory synapses^48,49^. Finally, we also implemented an experimentally observed^21–23^ anti-Hebbian rule in the second inhibitory population (Fig. 3C). Unlike its Hebbian counterpart, the anti-Hebbian rule leads to indefinite increases in the firing rate of the postsynaptic neuron, because correlated activity decreases synaptic weights (only sole presynaptic spikes increase synaptic weights; Methods). The anti-Hebbian plasticity rule is thus unstable (Fig. 3C, middle). We found that we could prevent catastrophe without incorporating additional, complex dynamics by using a variable learning rate for the anti-Hebbian rule. For simplicity, we decreased the learning rate exponentially over time (Fig. 3C, right), but this could also be achieved through top-down control (see Discussion).

**FIG. 3.**
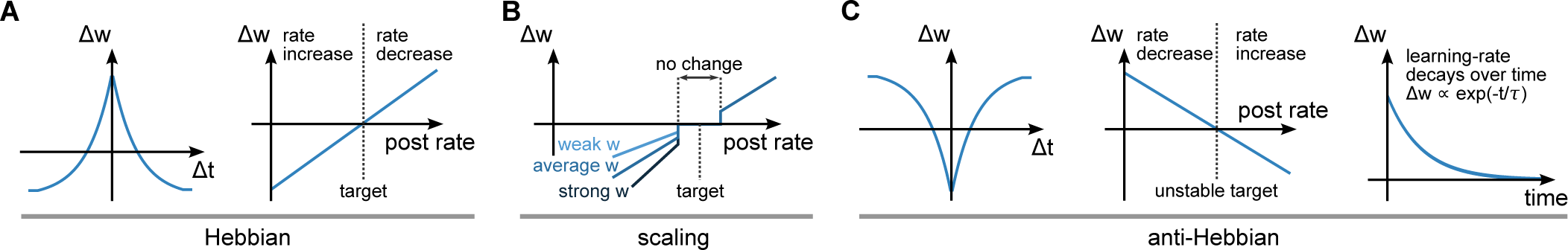
Synaptic plasticity models. **A**, Hebbian plasticity rule. *Left*, Spike-timing dependency; ∆*w* indicates level of synaptic change, and ∆*t* indicates interval between pre- and postsynaptic spikes. Coincident pre- and postsynaptic spikes elicit positive changes while presynaptic spikes alone elicit negative changes in synaptic strength^6^. *Right*, Synaptic changes (∆*w*) as a function of postsynaptic firing-rate. When the postsynaptic neuron’s firing-rate is above the target rate, inhibitory synapses increase in weight and, as a consequence, the postsynaptic neuron’s firing-rate decreases. The opposite happens for when the postsynaptic neuron’s firing-rate is lower than the target rate^6^ (Methods). **B**, Synaptic scaling rule. Changes in synaptic strength (∆*w*) as a function of the postsynaptic neuron’s firing-rate. When the postsynaptic neuron’s firing-rate is lower than a *lower bound* threshold, inhibitory synapses decrease, proportionally to their current strength. When the postsynaptic neuron’s firing-rate is higher than a *upper bound* threshold, inhibitory synapses increase. Because of the lower and upper bounds, there is a region with no change around the target rate. **C**, Anti-Hebbian plasticity rule. *Left*, Spike-timing dependency. Presynaptic spikes elicit positive changes, while coincident pre- and postsynaptic spikes elicit negative changes in synaptic weights. *Middle*, Changes in synaptic efficacy (∆*w*) as a function of the postsynaptic firing-rate. The target rate of anti-Hebbian plasticity rule is unstable. *Right*, Evolution of the learning-rate of the anti-Hebbian plasticity model. Due to its unstable nature, we set the learning-rate to decay exponentially over time.

### Shaping and modulating a single inhibitory population

To begin, we constructed a standard cortical circuit motif with one excitatory and one inhibitory population^6,18,50–52^ (Fig. 4A, top). We followed previous work showing that the Hebbian plasticity rule (Fig. 2A,B) changes inhibitory synapses to provide precisely balanced inputs^6^, such that both excitatory and inhibitory weight profiles are shaped according to previous experimental observations^12^ (Fig. 4A, bottom). Afferent synaptic weights were set so to allow average post-synaptic firing rates of approximately 5 Hz for natural inputs (Fig. 4B). We then changed the gain of all inhibitory afferents by modulating their firing rates, from 50% to 150% of control rates. This change of input balance translated into changes in output rates (Fig. 4C, bottom), and spike patterns (Fig. 4B, middle and right). When inhibition was equal or larger than excitation, the output was largely uncorrelated to any given input signal (Fig. 4D, top). When inhibitory firing rates fell below 90% of the control condition, the output first began to correlate with the preferred input signal. When inhibition became even weaker, the correlations increased, and even non-preferred signals were articulated in the postsynaptic firing patterns (Fig. 4D, bottom).

**FIG. 4.**
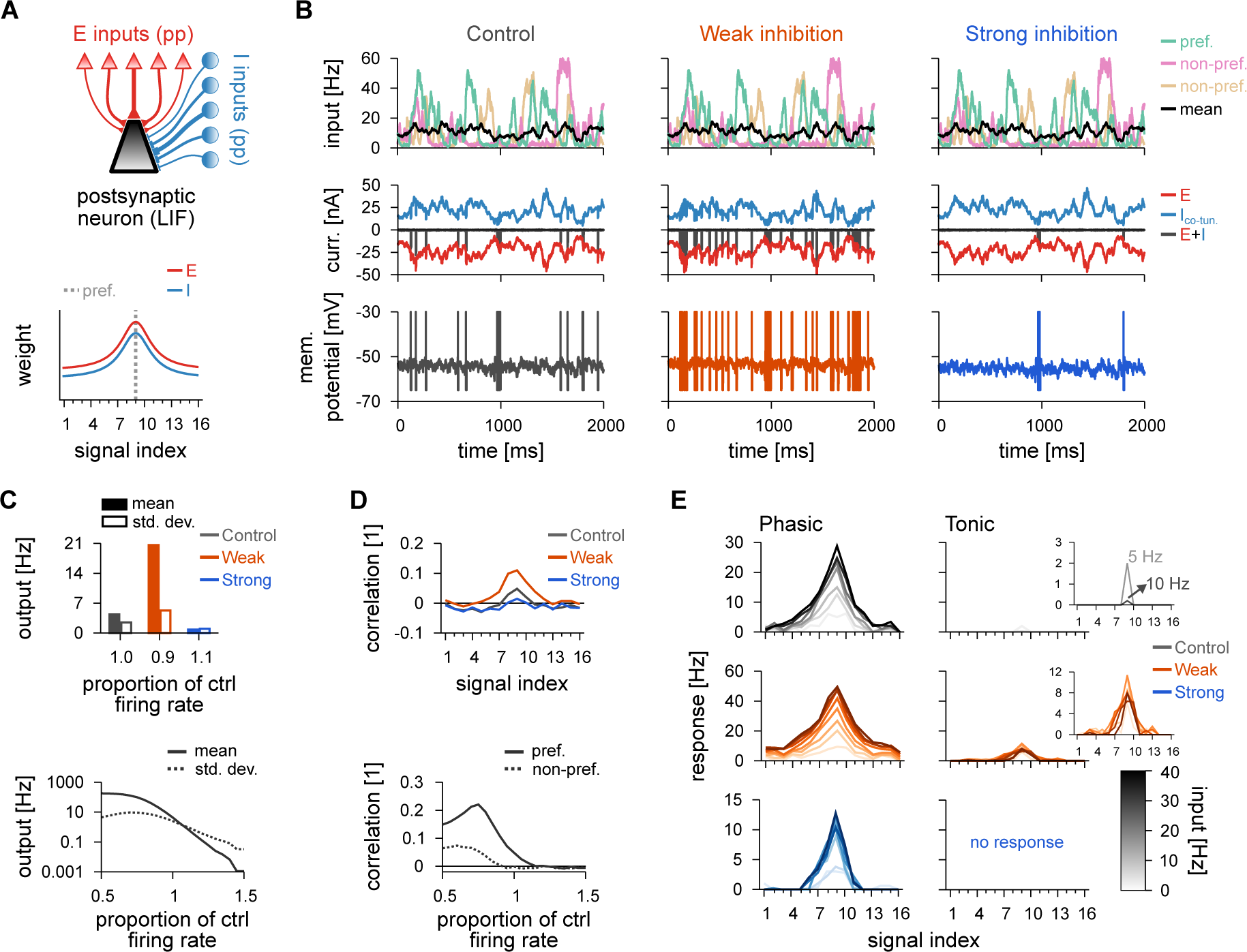
Postsynaptic response for a model with a single inhibitory population. **A**, Schematic of the circuit with a single inhibitory population (top). Pre-synaptic spikes were generated as point-processes (pp), for both excitatory (red; 16 signals) and inhibitory (blue; 16 signals) inputs, and fed into a single-compartment leaky integrate-and-fire neuron (LIF). Schematic of the synaptic weight profiles (bottom). Average weight (y-axis) for different input signals (x-axis); preferred signal is pathway no. 9 (grey dashed line). **B**, Average firing-rate of the preferred, and two non-preferred inputs and mean of all inputs (top row), excitatory and inhibitory input currents (middle row), and membrane potentials (bottom row), for control (left), decreased (middle) and increased (right) inhibition. Control case is hand-tuned for postsynaptic firing-rates of ~5 Hz. Decreased (increased) inhibition lowered (raised) inhibitory firing-rates by 10%, respectively. **C**, Average and standard deviation of the postsynaptic firing-rate in response to natural input for the three explored cases (top), and as a function of the inhibitory firing-rate (bottom). **D**, Pearson correlation between postsynaptic firing-rate and excitatory input firing-rates for different input signals for the three conditions in B (top). Correlation between output activity and preferred (continuous line) or non-preferred (dashed line) inputs as a function of the inhibitory firing-rate (bottom). **E**, Response to a pulse input in the phasic (left; first 50 ms), and tonic (right; last 50 ms) periods. Firing rate computed as the average number of spikes (for 100 trials) normalised by the bin size (50 ms). Each line corresponds to a different input strength; from light (low amplitude pulse) to dark (high amplitude pulse) colours. Insets show tonic response for control and decreased inhibitory firing-rates.

Transient presynaptic activity pulses caused strong phasic responses in the balance state when they were delivered through the afferents of the preferred inputs (Fig. 4E, top row). Stimuli from non-preferred afferents were largely ignored. This discriminability between transients of low or high amplitude pulses decreased when inhibition was down-regulated (Fig. 4E, middle row) such that pulse stimuli from all signal groups caused a response. Increased inhibition, on the other hand, completely abolished transient responses to non-preferred afferents (Fig. 4E, bottom row). In all three cases (balanced control, weak and strong inhibition), the postsynaptic neuron elicited most of its spikes within the phasic period of the total 100 ms input step (Fig. 4E). This indicates that strong postsynaptic responses are mostly driven by the onset of the presynaptic stimulation rather than the stimulus being integrated slowly over time, consequence of the precise balance of excitatory and inhibitory inputs^6^.

Thus, a single inhibitory population, even with tuned weights, could not affect the postsynaptic receptive field via only the modulation of the inhibitory firing-rate. To test whether an additional inhibitory population would allow for more sophisticated control of postsynaptic activity, we constructed a model with different plasticity rules, which were applied to two different populations of inhibitory inputs.

### Plasticity shapes inhibitory weight profiles and receptive fields

To study how plasticity can shape the emergence of distinct synaptic weight profiles, we incorporated inhibitory synaptic plasticity mechanisms into a model with two inhibitory populations. We started with a symmetric Hebbian plasticity rule in one of the two inhibitory populations: coincident pre- and postsynaptic spikes potentiated synapses whereas sole presynaptic spikes depressed synapses^6^ (Fig. 3A). The synapses of the excitatory and the other inhibitory population remained fixed (Fig. S1). Simulations began with tuned excitatory synapses and flat inhibitory weight profiles in both inhibitory populations (Fig. S1A).

After 30 minutes of stimulation with natural inputs (cf. Fig. 2B), inhibitory weights of the plastic population stabilised (Fig. S1D-G). Whether the target firing rate (Fig. S1B,C) was reached depended on the synaptic strength of the other, static population of inhibitory synapses. If the static weights were weak, the plastic synapses increased their strength until the target firing rate was reached (Fig. S1C). If the static population provided strong inhibition (and thus kept postsynaptic firing below the target rate), weights from the plastic population would eventually vanish – before the target firingrate could be reached (Fig. S1C,G). Consequently, the shape of the static population determined the shape of the plastic population (Fig. S1D,E). As expected, the input/output correlation of the postsynaptic responses followed the effective synaptic weight profile (Fig. S3A, cf. Fig. S1E), with distinct input/output correlations for turning either of the populations off (Fig. S3B). The Hebbian plasticity rule, due to the strengthening of synapses for coincident pre- and postsynaptic spikes, thus complemented additional inhibitory synaptic connectivity in establishing a state of detailed balance of excitatory and inhibitory inputs.

#### Hebbian and scaling plasticity rules

Next, we introduced plasticity to the second population of inhibitory afferents. We tested two different rules, beginning with a homeostatic plasticity rule which (multiplicatively) scaled synapses down and additively potentiated synapses so that a fixed-point for the post-synaptic firing-rates was reached (Fig. 3B; Methods). With the homeostatic rule co-active, the *Hebbian* synapses – connections changing according to the Hebbian plasticity rule – developed a co-tuned profile from initially random weights (Fig. 5A, top; Fig. 5C, left), while the synapses following the scaling rule collapsed to a single value (Fig. 5A, bottom; Fig. 5C, right; see Methods for mathematical analysis). Consequently, the post-synaptic neuron received precisely balanced inputs (Fig. 5B). The two plasticity rules cooperate to impose an average postsynaptic activity, and thus naturally work in harmony.

**FIG. 5.**
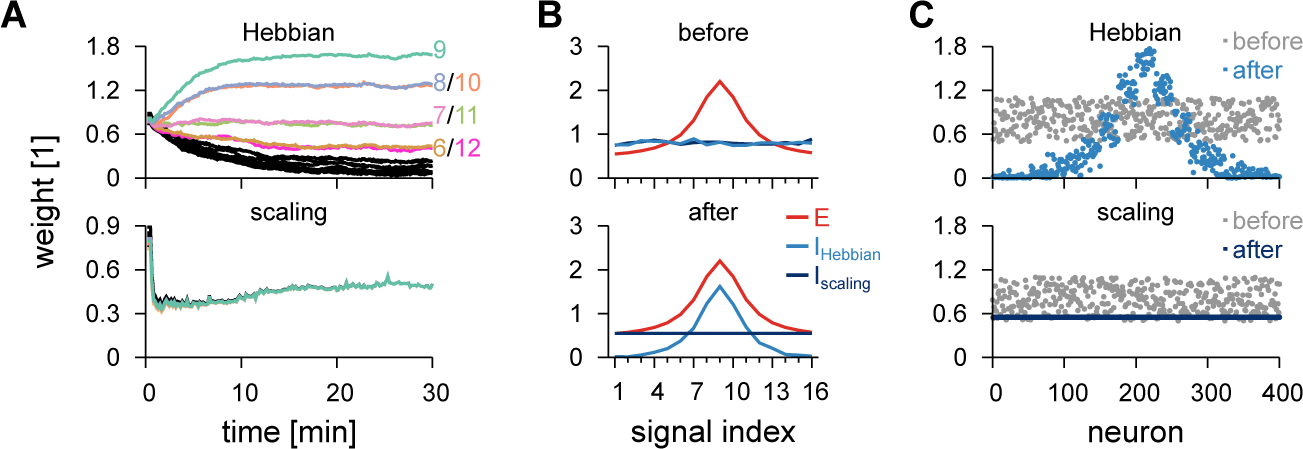
Simultaneous learning of two inhibitory profiles via Hebbian and homeostatic scaling plasticity rules. **A**, Temporal evolution of inhibitory synaptic weights when one inhibitory population follows a Hebbian plasticity rule (top) and the other population follows a synaptic scaling plasticity rule (bottom). **B**, Initial (top) and final (bottom) weight profiles from A with excitatory weights for reference. **C**, Individual synaptic weights before and after learning for synapses following the Hebbian plasticity rule (left) and synapses following the scaling plasticity rule (right).

We then studied the effects of differentially modulating the activity of the two inhibitory populations after their tuning curves had been established by the plasticity rules described above. First, we focused on the interaction of the connectivity created by Hebbian and the scaling plasticity rules (Fig. 6A, top; cf. Fig. 5), i.e., a *co-tuned* population and a *flat* population (Fig. 6A, bottom). We compared the output of the neuron in three scenarios: with both inhibitory populations active (control); with the co-tuned population inactive; and with the flat population inactive (Fig. 6B-E).

With both populations active, the input-output correlation was indistinguishable from a model with one, homogeneous inhibitory population (Fig. 6D, top; cf. Fig. 4D, top), because the two populations (co-tuned and flat) were mimicking the effect of the single (co-tuned) population. Deactivating either population (while increasing the firingrate of the other to maintain the same average output firing rate of 5 Hz in the modulated conditions, Fig. 6C) had pronounced effects on postsynaptic responses. Fluctuations in firing rate and membrane potential increased in both cases (Fig. 6B,C). When the co-tuned inhibitory population was turned off, the emerging imbalance of excitation and inhibition unmasked the excitatory tuning curve, thus increasing the chance of action potential generation when preferred signal populations were active (Fig. 6B, middle). The compensatory increase in the activity of the flat population further quenched non-preferred excitatory signals, leading to anti-correlated responses for non-preferred input signals (Fig. 6D, purple), reflecting the lack of postsynaptic firing during periods in which non-preferred signals were active (Fig. 6B, middle). The opposite effect could be observed when the flat population was deactivated. In this case, the lack of inhibition for non-preferred signals gave rise to input/output correlations for non-preferred signals, while preferred signals saw no response (Fig. 6B, right and Fig. 6D, green).

**FIG. 6.**
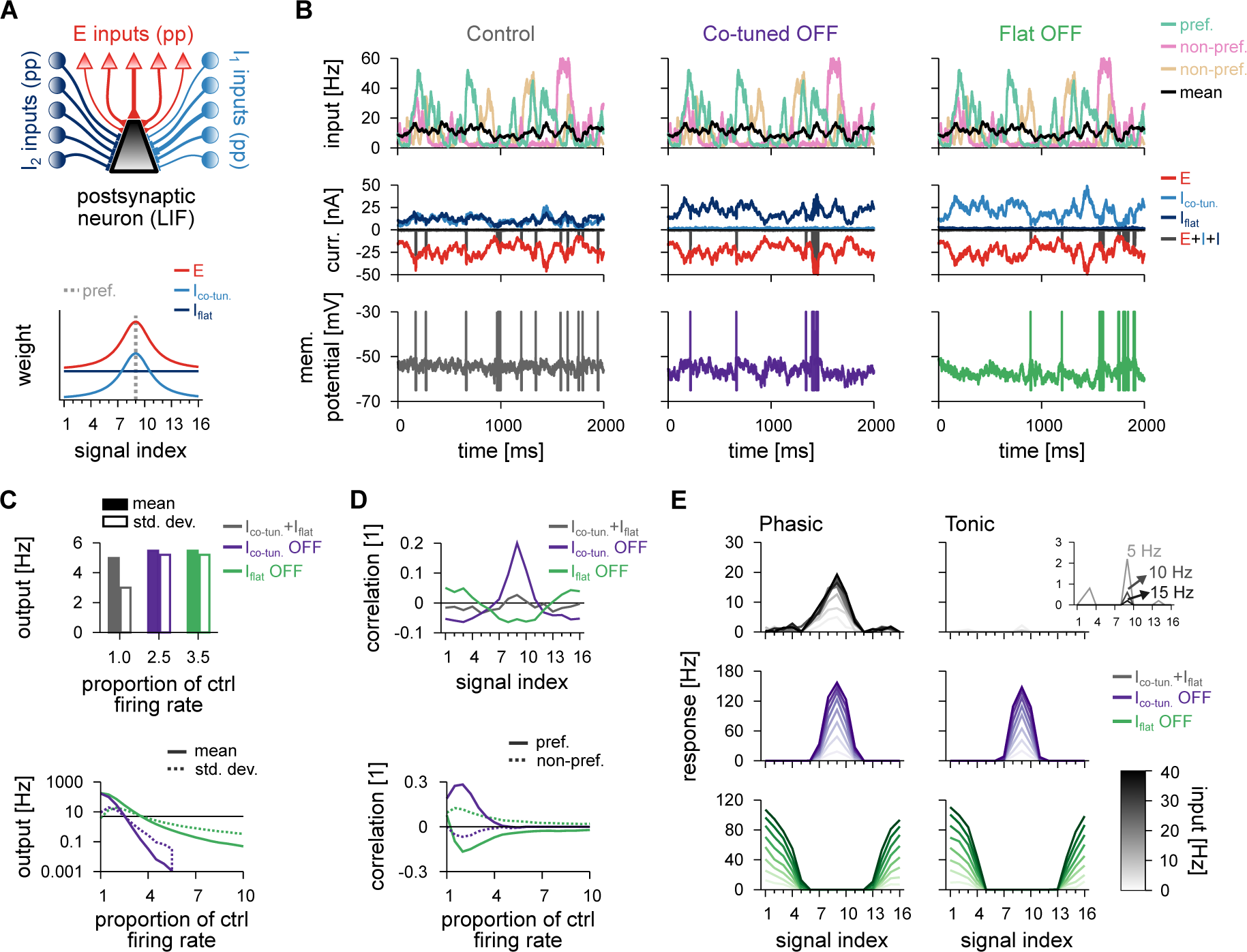
Postsynaptic response for the model with co-tuned and flat inhibitory populations. **A**, Schematic of the circuit with two inhibitory populations (top); *I*_1_ corresponds to the co-tuned population and *I*_2_ to the flat population. Pre-synaptic spikes were generated as point-processes (pp) and fed into an LIF. Schematic of the synaptic weight profile (bottom). Average weight (y-axis) for different input signals (x-axis); preferred signal is pathway no. 9 (grey dashed line). **B**, Average firing-rate of the preferred and two non-preferred inputs and mean of all inputs (top row), total excitatory current and inhibitory currents of both populations (middle row), and membrane potential (bottom row), for control (left), co-tuned (middle) and flat (right) population inactive. **C**, Average and standard deviation of the postsynaptic firing-rate in response to natural input for the three cases (top), and as a function of the inhibitory firing-rate (bottom). **D**, Pearson correlation between postsynaptic firing-rate and excitatory input firing-rates for different input signals for the three conditions in B. Correlation between output activity and preferred (continuous line) or non-preferred (dashed line) as a function of the inhibitory firing-rate of each inhibitory population (bottom). **E**, Response to a pulse input in the phasic (left; first 50 ms), and tonic (right; last 50 ms) periods. Firing rate computed as the average number of spikes (for 100 trials) normalised by bin size (50 ms). Each line corresponds to a different input strength; from light (low amplitude pulse) to dark (high amplitude pulse) colours. Insets show tonic response for control firing-rates.

Transient responses, when compared to the unmodulated control case (Fig. 6E, top), were substantially increased for preferred inputs when the co-tuned population was deactivated, and the response to non-preferred signals was completely diminished (Fig. 6E, middle). When the flat population was deactivated, the postsynaptic neuron responded strongly to the non-preferred inputs, but not to preferred inputs (Fig. 6E, bottom). Interestingly, modulating either of the inhibitory populations had similar effects on the postsynaptic response both in phasic and tonic periods, in contrast with the unmodulated control case, in which only phasic responses were postsynaptically elicited (Fig. 6E). Again, this reflects the state of balance between excitation and inhibition in the unmodulated control case, which only reveals transient input dynamics.

#### Hebbian and anti-Hebbian plasticity rules

Instead of a purely homeostatic scaling rule, we also tried an experimentally observed^21–23^ anti-Hebbian rule in the second inhibitory population (Fig. 3C). The anti-Hebbian rule, unlike the Hebbian, decreases synaptic weights for correlated activity, and sole presynaptic spikes increase synaptic weights. Such a rule can only either indefinitely increase the firing rate of the postsynaptic neuron or decrease it to zero (Methods). We accounted for the unstable nature of the anti-Hebbian plasticity rule (Fig. 3C, middle) by controlling its learning rate, such that it decreased exponentially over time (Fig. 3C, right). With both Hebbian and anti-Hebbian rules active, initially random weights evolved into co-tuned and counter-tuned synaptic weight profiles (Fig. 7). As learning slowed down due to the decreasing learning rate, the *anti-Hebbian* synapses – connections changing according to the anti-Hebbian plasticity rule – stabilised, and Hebbian synapses ceased to change once the target firing rate was reached (Fig. 7B,C).

**FIG. 7.**
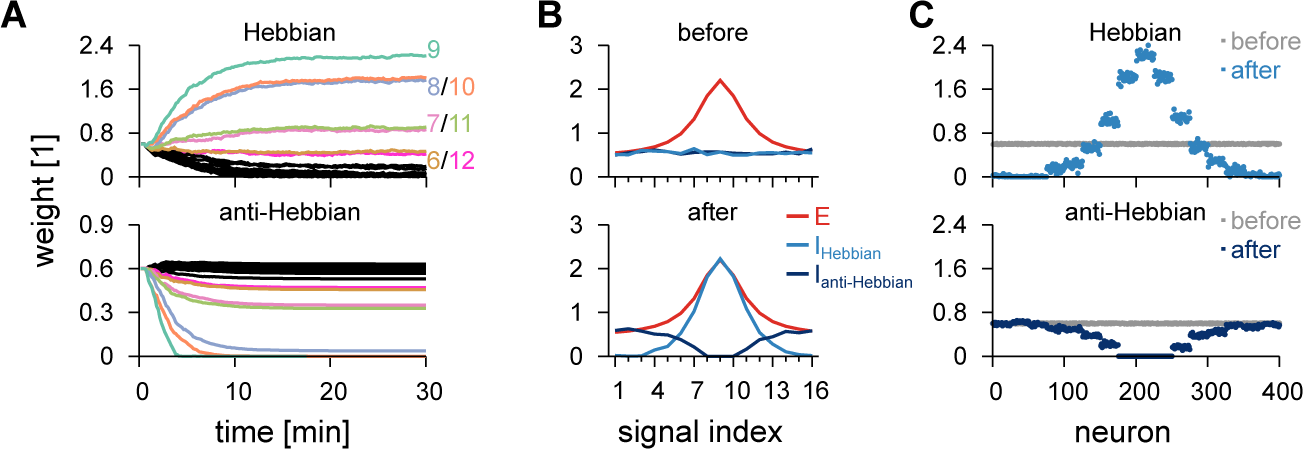
Simultaneous learning of two inhibitory profiles via Hebbian and anti-Hebbian plasticity rules. **A**, Temporal evolution of inhibitory synaptic weights when one inhibitory population follows a Hebbian plasticity rule (top) and the other population follows an anti-Hebbian plasticity rule (bottom). **B**, Initial (top) and final (bottom) weight profiles from A with excitatory weights for reference. **C**, Individual synaptic weights before and after learning for synapses following the Hebbian plasticity rule (left) and synapses following the anti-Hebbian plasticity rule (right).

Postsynaptic dynamics with two inhibitory populations with tuning that resulted from the combination of the Hebbian and the anti-Hebbian plasticity rules (Fig. S2A), i.e., *co-tuned* and *counter-tuned* populations, were similar to that with cotuned and flat inhibitory populations. In the unmodulated balanced state, output behaviour is near identical to previous results (Fig. S2B-D, control). The main distinction between the models with counter-tuned or flat inhibitory profiles is how they complemented the co-tuned inhibitory currents: the flat inhibition produced currents that tracked the co-tuned inhibitory currents, whereas counter-tuned inhibition produced inhibitory currents that were largely uncorrelated to the co-tuned inhibitory currents (Fig. S2B, left; compare with Fig. 6B, left).

When either the co- or the counter-tuned inhibitory populations were inactivated, fluctuations in both firing rate and membrane potential increased considerably (Fig. S2B, middle and right). Deactivation of the co-tuned population resulted in positive correlation between postsynaptic activity and preferred signals, and negative correlation between output and non-preferred signals (Fig. S2D, purple). For transient stimulation, there was no discernible difference to the model with flat inhibition in the control state (Fig. S2E, top).

Turning off counter-tuned inhibition (Fig. S2B-E) also had similar results in the postsynaptic response as turning off the flat inhibition (cf. Fig. 6B-E), i.e., non-preferred input produced output activity with positive correlation (Fig. S2D) and strong postsynaptic activity for transient activation (Fig. S2E, bottom). Unlike before, turning off co-tuned inhibition produced elevated firing-rate responses also for transient stimuli from signals directly neighbouring the preferred input (Fig. S2E, middle row, compare with Fig. 6E, middle).

### Quantitative differences of inhibitory profiles

For a better understanding of the differences between the three conditions studied here (one inhibitory population, co-tuned & flat and co- & counter-tuned) we compared different modulation schemes quantitatively. We introduced the parameter ∆*C* = 0.5(*C*_pref_ − *C*_non-pref_), i.e., 50% of the difference in input/output correlation between preferred, *C*_pref_, and nonpreferred, *C*_non-pref_, signals (Methods). Ideally, the sensory system should present three distinct responses for the three different modulatory conditions, which are captured by different values of ∆*C*. With unmodulated input (control), the output neuron should present correlated activity with all input groups, and thus ∆*C* ≈ 0. Modulated inputs (by decreasing the activity of either of the inhibitory populations) should correlate preferred (for one inhibitory inactive) and non-preferred (for the other population inactive) to the output activity. This results in ∆*C* > 0 for correlated output/preferred signals, and ∆*C* < 0 for correlated output/non-preferred signals.

In the control condition, we observed similar ∆*C* ≈ 0 in all models (Fig. 8A, grey), reflecting low levels of correlation between output and input signals (Fig. 8B, top). With down-regulated inhibition, ∆*C* increased slightly in the model with one homogeneous inhibitory population. ∆*C* increased more considerably in a two-population model in which the cotuned population was inactive (Fig. 8A, purple), confirming an increased correlation between preferred signal and output (Fig. 8B, middle). When the flat or the counter-tuned inhibitory populations were inactivated, we observed postsynaptic responses even to non-preferred input signals (Fig. 8B, bottom), which led to negative ∆*C*. Inactivating the counter-tuned inhibition resulted in a slightly better discrimination (larger negative ∆*C*) of non-preferred input signals (Fig. 8A, green).

**FIG. 8.**
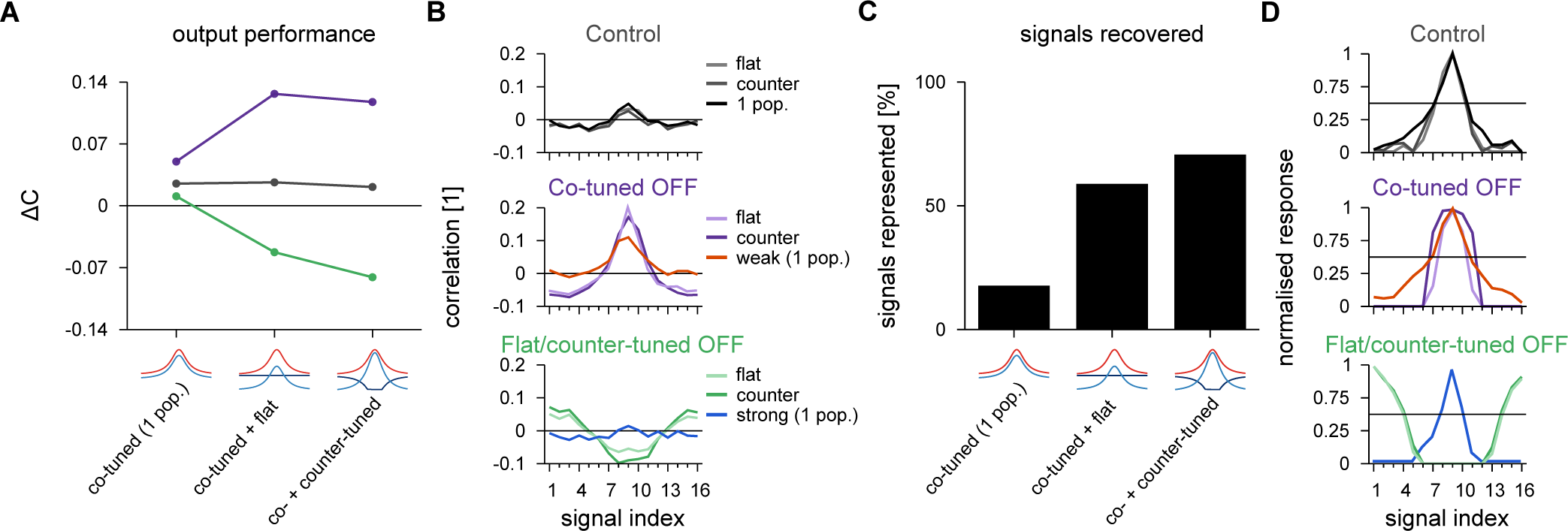
Comparison of postsynaptic responses receiving co-tuned & flat or co-tuned & counter-tuned inhibitory populations. **A**, Performance index as the difference in input/output correlation between preferred and non-preferred signal. Ideal outcome is ∆*C* = 0 for control case (grey), ∆*C* > 0 for co-tuned population inactive (purple), and ∆*C* < 0 for flat or counter-tuned populations inactive (green). We added the values for a single inhibitory population with control (grey), weak (purple) and strong (green) inhibitory inputs for comparison. **B**, Pearson correlation between postsynaptic firing-rate and excitatory input firing-rates for different signal indices from Figs. 4D, 6D and S2D, replotted for reference. **C**, Signals recovered in the pulse input paradigm. Signals represented are calculated as the percentage of signal afferents that activate the postsynaptic neuron with more that half the spikes of the maximum response for the three cases considered for the circuits analysed together. **D**, Normalised phasic response to a pulse input of 40 Hz, from Figs. 4E, 6E and S2E, replotted for reference. Horizontal line indicates 50% of maximum response.

To compare pulse responses of the three models, we quantified which input signal groups elicited a substantial response to a pulse signal. We defined the number of *signals recovered* (Fig. 8C) as the number of responses with more than 50% of the maximum postsynaptic firing-rate (Fig. 8D). The single inhibitory population model could only produce responses to preferred input signals, while co-modulation of two inhibitory populations could promote responses to non-preferred input signals, as well. Counter-tuned population achieved better (i.e., broader) postsynaptic control than flat inhibition (Fig. 8C).

The addition of a second population of inhibitory inputs thus gives rise to a more flexible response to varying stimuli. In summary, our results shed light on the role of the many types of interneurons in cortical areas^1,2,4^, and show the benefits of combining different biologically inspired plasticity rules in neuronal networks.

## Discussion

We investigated how several distinctly tuned inhibitory connectivity profiles emerge through biologically reasonable plasticity rules and how they interact with a tuned excitatory connectivity profile in a receptive field-like paradigm. We found that the two aspects of selective attention – enhancing response to targets, and suppressing the response to distractors – were implemented in our model by two types of disinhibition. Our results indicate a simple neuronal mechanism to help disentangle (or bind) parallel sensory input streams and may represent a step towards understanding the neural basis of intricate behaviours such as the *cocktail party e*ff*ect* – focusing on a single voice in a crowded, cacophonous place.

### Modulation of receptive field response

Our findings also fit well with recent experimental results showing that pyramidal neurons in sensory areas of the cortex change their response to external stimuli depending on the context of the signal or attentional state^29–33^. For example, principal neurons in macaque V4 respond to monochrome images of varying hues with variable response amplitude that is consistent with specific colour-tuning. However, the preferred colour response of the neurons changes when naturally coloured images are shown. In macaque V1, principal neurons can change the preferred orientation of visual stimuli when a pure tone is played alongside the visual stimulation^30^. In the framework of our model, such a change in preference could be explained with differential input to the two inhibitory populations, or by changes in their gains through contextual neuromodulation. Similarly, up to 20% of neurons in all areas of the mouse visual system^34^ were recently shown to change their preferred orientation according to the (spatial and temporal) frequency of the drifting gratings used in the experiments. These effects could also be explained by temporal fluctuations in the interaction of the two inhibitory populations, and the concurrent changes in transient responses of our model.

### Neuron types

The architecture of our model maps easily onto the neocortical microcircuit^1,4^. Co-tuned inhibition, e.g., may originate from parvalbumin-positive (PV+) interneurons. As the main source of inhibition to pyramidal cells, PV+ interneurons target postsynaptic neurons with similar preferred orientation^35^, and activation of these neurons leads to broadened selectivity^35^ (but see Lee *et al.* ^53^). Flat or counter-tuned inhibition may arrive from somatostatin-positive (SOM+) interneurons with their less selective connectivity patterns^35^. This interpretation is also in line with recent evidence suggesting that top-down visual attention relies on local inhibitory circuitry in primary visual cortex^54^. In this scheme, PV+ and SOM+ neurons inhibit pyramidal cells, while vasoactive intestinal peptide-positive (VIP+) neurons suppress other inhibitory interneurons, acting as a source of disinhibition. Direct manipulation of SOM+, PV+ and VIP+ neurons confirms these respective roles in inhibition and disinhibition in both visual^55^ and auditory cortices^14,36^. Additionally, Zhou *et al.* ^54^ reported that VIP+ neurons received excitatory top-down inputs from the rodent cingulate cortex, leading to a narrow selectivity profile of pyramidal cells when cingulate inputs are active active and broad tuning when cingulate cortex is silent. Finally, blocking cortical inhibition reduces the stimulus-selectivity of cortical neurons^56,57^ (but see Nelson *et al.* ^58^).

### Balance between excitatory and inhibitory inputs

In our model, we aimed for precise balance of excitation and inhibition, by way of a Hebbian-like inhibitory plasticity rule^6^, and accordant with evidence of excitatory and inhibitory co-tuning in cat visual cortex^59^, rodent auditory cortex^12,54,60^ and rodent hippocampus^61^, and temporal correlations in neighbouring excitatory and inhibitory synapses^62^. Consistent with earlier work, we could modulate the efficacy of a single inhibitory population to enhance the output correlation with the preferred input^18,19^, but the flexibility of the control mechanism was very limited and non-preferred signals never evoked faithful responses.

### Inhibitory synaptic plasticity

To explore how different inhibitory synaptic populations could form and interact, we split the inhibitory afferents into two populations and implemented a Hebbian-like inhibitory plasticity rule^6,20^ in one population that was co-active with either a homeostatic scaling^48,49^ or an anti-Hebbian^21–23^ plasticity rule. The scaling plasticity rule acted locally, but squeezed the distribution of all synaptic strengths to a narrow regime, providing a parsimonious explanation for the un-tuned, *blanket* inhibition often encountered in experiments^63,64^, and providing easy means for modulating postsynaptic responses independently of the presynaptically-tuned weight profiles. The anti-Hebbian rule was naturally unstable, i.e., it could lead to infinite strengthening of weights and thus silent networks. Our implementation reinforces this outcome because inhibitory inputs are always active.

It is unclear how biological circuits would avoid such catastrophe, but in our model we could balance the effect of the two opposing rules and remain at plausible levels of postsynaptic activity by including a modulatory term that controlled the learning rate of the anti-Hebbian plasticity rule. While this mimics some of the observed modulatory control of plasticity through other neuronal types^12,65–67^, the reality is likely more complex, and possibly relies on finely orchestrated interaction of several different plasticity rules^9,68^. Additionally, if the inhibitory neurons are driven laterally by excitatory neurons that lack excitatory recurrence, a form of anti-Hebbian plasticity is also stable^24^. No matter what form the ultimate mechanism may take, it is unlikely that it will affect the generality of our results.

### Parallels to artificial neural networks

Interestingly, artificial networks have been shown to develop similar receptive field profiles to the ones explored here when they are trained to solve multiple tasks^69^. Yang *et al.* ^69^ have shown that clusters of neurons can acquire co-tuned or flat connectivity, which are controlled by context-encoding signals. These results hint at the possibility that biological and artificial systems may utilise similar strategies to solve context-dependent filtering tasks.

### Additional biological complexity

To explore the interaction between two distinct inhibitory plasticity rules without confounds, we made the simplifying assumption that excitatory synapses would remain fixed (but see Clopath *et al.* ^51^, Litwin-Kumar and Doiron ^70^, and Zenke *et al.* ^68^). Obviously, inhibitory plasticity rules do interact with multiple additional rules and constraints like, e.g., excitatory or modulatory synaptic plasticity. Similarly, our model only considered a single postsynaptic neuron, with no feedback or lateral connectivity, which is thought to play an important role in cortical feature selectivity^71^, and was theoretically shown to provide the means for multiplicative and additive modulation of receptive fields, and surround suppression^72^. Finally, other possible functions beyond simple input filtering, such as multiplexing or amplifying temporally varying signal streams^73,74^, and one-shot learning^11^ must be considered. Our work only lays the groundwork for studies of multiple distinct plasticity rules in larger networks, with more complex excitatory-inhibitory interaction^9,75,76^

### Conclusion

We predict that various GABAergic interneurons in the same cortical region must obey a range of different inhibitory synaptic plasticity rules, to restore or reverse neuronal stimulus-selectivity as appropriate and necessary. Such evidence would inform the theoretical framework presented here, and in turn inspire future computational modelling.

## Materials and Methods

Detailed methods can be found below (after references).

## Software and code availability

Simulations were run in Fortran, compiled with Intel Fortran Compiler 19.0 on an Intel-based Linux computer (Debian 9; i9-9900X processor; 32 GB memory). Codes will be made available online upon publication^77^. Individual plots were generated with Gnuplot. Figures were generated with Inkscape.

## Acknowledgements

We thank the members of the Vogels group, and particularly Georgia Christodoulou and William Podlaski, for fruitful discussions. This work was supported by a Research Project Grant by the Leverhulme Trust (RPG-2016-446; EJA), a Sir Henry Dale Fellowship by the Wellcome Trust and the Royal Society (WT100000; EJA, TPV), a Wellcome Trust Senior Research Fellowship (214316/Z/18/Z; EJA, TPV), an ERC Consolidator Grant (SYNAPSEEK; EJA, TPV), and a Claredon Scholarship from the University of Oxford (AIL).

## Author contributions

EJA and TPV designed research; EJA and AIL carried out the simulations and analysis; EJA, AIL and TPV wrote the manuscript.

## Competing interests

The authors declare no competing financial interests.

## Methods

Codes for all results are openly available at GitHub, repository https://github.com/ejagnes/attentional_switch.

### Neuron model

To investigate changes in neuronal response due to specific inhibitory connectivity motif we simulated a postsynaptic leaky integrate-and-fire neuron (LIF) receiving excitatory and inhibitory afferents. Postsynaptic neuronal membrane potential dynamics is governed by

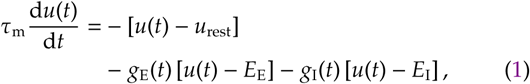

where *u*(*t*) is the somatic voltage at time *t*, τ_m_ = *RC* is the membrane time constant (membrane resistance, *R*, times membrane conductance, *C*), *u*_rest_ is the resting membrane potential, and *E*_E_ and *E*_I_ are the reversal potential for excitatory and inhibitory synapses, respectively. Synaptic conductances, *g*_E_(*t*) and *g*_I_(*t*), evolve according to

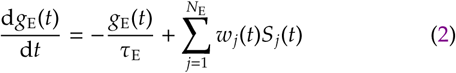

and

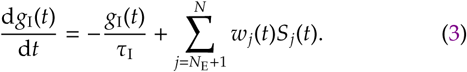

Both excitatory and inhibitory conductances decay exponentially to zero with time constants τ_E_ and τ_I_, respectively. Presynaptic action potentials trigger increase in synaptic conductances through the sum of Dirac delta functions,

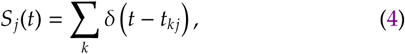

where *t*_*kj*_ is the time of the *k*th spike of presynaptic afferent *j*. The contribution of a given presynaptic afferent *j* to changes in conductances is given by the synaptic weight *w*_*j*_(*t*), which was fixed for excitatory synapses and could change over time due to plasticity mechanisms for inhibitory synapses. The total number of presynaptic afferents is *N* = *N*_E_ + *N*_I_, with *N*_E_ being the number of excitatory and *N*_I_ of inhibitory presynaptic afferents.

An action potential is triggered at the postsynaptic neuron once its membrane potential *u*(*t*) crosses the spiking threshold *u*_th_ from below. The membrane potential is then instantaneously reset to *u*_reset_, being clamped at this value for the duration of the refractory period, τ_ref_. The postsynaptic spike train is described here as a sum of Dirac deltas,

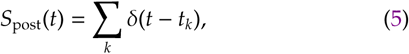

where *t*_*k*_ is the time of the *k*th spike of the postsynaptic neuron, or the time the membrane potential crosses the spiking threshold from below. Parameters used for the postsynaptic neuron are detailed in Table I.

**TABLE I.** Simulation parameters for postsynaptic neuron.

| Parameter | Symbol | Value | Figs. |
| --- | --- | --- | --- |
| Membrane time constant | $\tau_m$ | 30 ms | 4 – 8, S1 – S3 |
| Resting potential | $u_{\text{rest}}$ | –65 mV | 4 – 8, S1 – S3 |
| Excitatory reversal potential | $E_E$ | 0 mV | 4 – 8, S1 – S3 |
| Inhibitory reversal potential | $E_I$ | –80 mV | 4 – 8, S1 – S3 |
| Excitatory time constant | $\tau_E$ | 5 ms | 4 – 8, S1 – S3 |
| Inhibitory time constant | $\tau_I$ | 10 ms | 4 – 8, S1 – S3 |
| Spiking threshold | $u_{\text{th}}$ | –50 mV | 4 – 8, S1 – S3 |
| Reset potential | $u_{\text{reset}}$ | –60 mV | 4 – 8, S1 – S3 |
| Refractory period | $\tau_{\text{ref}}$ | 5 ms | 4 – 8, S1 – S3 |
| Simulation time step | $\Delta t$ | 0.1 ms | 4 – 8, S1 – S3 |

### Inputs

To mimic experimentally observed synaptic input profiles^12^, we divided the synaptic inputs into *P* signal groups (*µ* = 1, …, *P*) that share the same fluctuation in firing rate. We tested two cases: (*i*), natural input, and (*ii*), pulse input. Both are described below.

#### Natural input

For presynaptic activity mimicking a natural input, activity follows an inhomogeneous Poisson process that changes according to a modified Ornstein-Uhlenbeck (OU) process. We first defined an auxiliary variable for each pattern, *y*_*µ*_(*t*), that follows a stochastic first-order differential equation given by

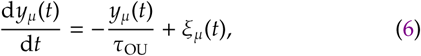

where *µ* is the signal group index, τ_OU_ is the time constant for the decaying process that changes due to a Gaussian noise term ξ_*µ*_(*t*) with unitary standard deviation. The mean value of the variable *y*_*µ*_ is zero, and thus it assumes positive and negative values with same probability (for long periods).

The spike train of an afferent in a given signal group *µ* is given by the variable *ν*_*Xµ*_(*t*) which is a rectified version of the auxiliary variable plus a term to generate background firing rate, *ν*_*X*bg_, where *X* indicates the presynaptic population; *X* = E for excitatory and *X* = I for inhibitory. The spike trains of the afferents of signal group *µ* are generated by

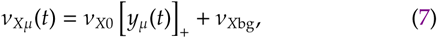

where *ν*_*X*0_ is the amplitude of the modulated firing rate fluctuations, and [⋅]_+_ is a rectifying function,

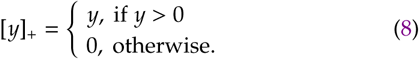

Note that due to the symmetry of *y*_*µ*_(*t*), an afferent is half the time in the background state and half the time in the active state.

Presynaptic action potentials were generated as an inhomogenous Poisson process according to the modified OU process described above and a fixed background firing rate. Additionally, we implemented a refractory period, τ_Eref_ for excitatory and τ_Iref_ for inhibitory inputs. Given the time step of the simulation ∆*t*, spikes of a presynaptic afferent that is part of the signal group *µ* are generated with a probability *p*_*Xµ*_(*t*) = *ν*_*Xµ*_(*t*)∆*t* if there was no spike elicited during the refractory period beforehand, and thus the average firing rate of a *X* = E (excitatory) or *X* = I (inhibitory) afferent that is part of the signal group *µ* becomes

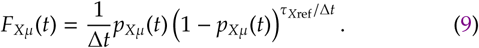

#### Pulse input

To test transient responses to brief changes in presynaptic activity we also quantified postsynaptic responses to pulse inputs. In this case, we simulated the postsynaptic neuron receiving inputs with constant background firing-rate. For 100 ms we increased the probability of presynaptic spikes for a given signal group *µ* by a factor *k*ν**^∗^, with *k* being an integer larger or equal than zero, and *ν*^∗^ = 5 Hz. Thus presynaptic spikes are generated by

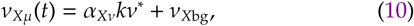

during the 100 ms step and by

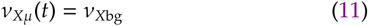

during only background activity. Parameter *α*_*X*ν**_ is a scalar that sets the ratio of excitatory and inhibitory firing rate.

Responses to the pulse input were divided in two bins: *phasic* and *tonic*. Phasic responses were defined as the postsynaptic activity elicited in the first 50 ms of the pulse input. Tonic activity was correspondingly defined as having occurred in the last 50 ms of the stimulus. We simulated 100 trials per input strength *k*ν**^∗^, and defined the response (for both phasic and tonic) as the average number of spikes on the period for the strength *k*ν**^∗^ minus the average number of spikes on the same period without extra input, multiplied by 20 to convert to Hz. We subtracted background spikes to ascertain that we quantified the response to the extra step input alone. Parameters used for inputs are detailed in Table II.

**TABLE II.** Simulation parameters for inputs.

| Parameter | Symbol | Value | Figs. |
| --- | --- | --- | --- |
| Number of excitatory afferents | $N_E$ | 3200 | 4 – 8, S1 – S3 |
| Number of inhibitory afferents | $N_I$ | 800 | 4 – 8, S1 – S3 |
| Number of signal groups | $P$ | 16 | 4 – 8, S1 – S3 |
| Refractory period for excitatory afferents | $\tau_{\text{Eref}}$ | 5 ms | 2, 4 – 8, S1 – S3 |
| Refractory period for inhibitory afferents | $\tau_{\text{Iref}}$ | 2.5 ms | 2, 4 – 8, S1 – S3 |
| Ornstein-Uhlenbeck process (OU) time constant | $\tau_{\text{OU}}$ | 50 ms | 2, 4 – 8, S1 – S3 |
| OU update time step | $\Delta T$ | 1 ms | 2, 4 – 8, S1 – S3 |
| Excitatory firing rate amplitude for OU | $v_{E0}$ | 5 Hz | 2, 4 – 8, S1 – S3 |
| Inhibitory firing rate amplitude for OU | $v_{I0}$ | 10 Hz | 2, 4 – 8, S1 – S3 |
| Excitatory background firing-rate | $v_{\text{Ebg}}$ | 2 Hz | 2, 4 – 8, S1 – S3 |
| Inhibitory background firing-rate | $v_{\text{Ibg}}$ | 4 Hz | 2, 4 – 8, S1 – S3 |
| Pulse amplitude reference | $v^*$ | 5 Hz | 4, 6, 8, S2 |
| Excitatory ratio for pulse input | $\alpha_{Ev}$ | 1 | 4, 6, 8, S2 |
| Inhibitory ratio for pulse input | $\alpha_{Iv}$ | 2 | 4, 6, 8, S2 |
| Synaptic weight profile amplitude | $r_0$ | 4 | 4 – 8, S1, S3 |
| Synaptic weight profile slope | $b$ | 0.25 | 4 – 8, S1, S3 |
| Preferred pattern index | $\mu_0$ | 9 | 4 – 8, S1 – S3 |
| Synaptic weight profile power | $c$ | 2 | 4 – 8, S1 – S3 |
| Simulation time step | $\Delta t$ | 0.1 ms | 2, 4 – 8 – S1, S3 |

### Synaptic tuning

Based on Vogels *et al.* ^6^, we used a synaptic weight profiles given by

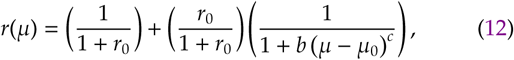

where *r*_0_, *b* and *c* are parameters defining the shape of the synaptic weight profile and *µ*_0_ defines the preferred signal group, which maximises *r*(*µ*); *r*(*µ*_0_) = 1. Note that *r*_0_ ≥ 1, *b* ≤ 1, *µ*_0_ > 0, and *c* is an even positive integer.

For simplicity we define ζ_*j*_ as the signal group that afferent *j* is part of. Thus excitatory synapses are set as

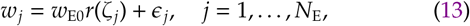

where *w*_E0_ is a normalisation factor for excitatory weights, and *ϵ*_*j*_ is a noise term drawn from a uniform random distribution between *ϵ*^∗^_E_ and *ϵ*^∗^_E_.

First we simulated a single inhibitory population with a tuned profile (Fig. 4), following Eq. 12 such that,

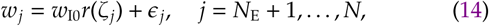

where *w*_I0_ is a normalisation factor for inhibitory weights, and *ϵ*_*j*_ is a noise term drawn from a uniform random distribution −between *ϵ*^∗^_I_ and *ϵ*^∗^_I_. The parameter *w*_I0_ was chosen so that a state of balance was enforced, with postsynaptic firing-rate of 5 Hz. Due to the small number of inhibitory afferents compared to the excitatory ones, and the difference in driving force, inhibitory weights were much larger than excitatory ones. Thus, to plot excitatory and inhibitory weights on the same scale we computed the *correcting* factor, *α*_*w*_, from

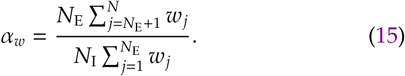

In all plots with excitatory and inhibitory weights, we plotted excitatory weights multiplied by the parameter *α*_*w*_.

Next, inhibitory afferents were divided in two types that connect to the postsynaptic neuron with different synaptic weight profiles. We started the inhibitory synaptic weights randomly and applied plasticity rules (details below). To compute input/output correlations and postsynaptic responses, we defined inhibitory weight profiles based on Eq. 12, i.e., we *hand-tuned* their shape according to the profiles learned due to the plasticity rules (details below).

We combined a co-tuned population with either a flat (Fig. 6) or a counter-tuned (Fig. S2) population. In both cases we based co-tuned weights on a modified version of Eq. 12,

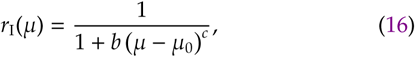

with *b*, *µ*_0_ and *c* the same as for Eq. 12. Inhibitory weights for the co-tuned profiles are chosen so that

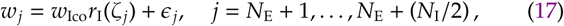

where 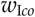 is a normalisation factor for inhibitory weights following the co-tuned synaptic weight profiles, and are different when combined with either the flat or the counter-tuned populations.

We set the flat population such that

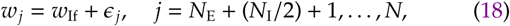

where *w*_If_ is the average for the flat population. The shape of the counter-tuned population was defined by

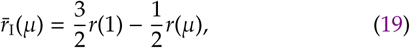

and synapses were hence tuned such that

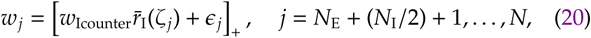

where [⋅]_+_ is a rectifier (Eq. 8), used to enforce only positive synaptic weights. The parameters 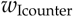 is a normalisation factor for the counter-tuned inhibitory populations.

When plasticity was simulated, initial conditions for all plastic inhibitory populations were flat with noise (Fig. 5, Fig. 7, Fig. S1 and Fig. S3),

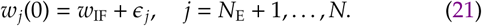

Parameters used for the tuning curves are detailed in Table II, and for synaptic weights in Table III. Both the average of the weights for flat population, *w*_If_, and the noise term, *ϵ*^∗^_I_, were distinct for different simulations.

**TABLE III.** Simulation parameters for weights.

| Parameter | Symbol | Value | Figs. |
| --- | --- | --- | --- |
| Excitatory baseline weight | $w_{E0}$ | 0.5 | 4 – 8, S1 – S3 |
| Noise parameter for excitatory weights | $\epsilon_E^*$ | 0.01 | 4 – 8, S1 – S3 |
| Inhibitory baseline weight (one inh. population) | $w_{I0}$ | 1.04 | 4, 8 |
| Inhibitory baseline weight (co-tuned & flat) | $w_{Ico}$ | 1.58 | 6, 8 |
| Inhibitory baseline weight (co-tuned & flat) | $w_{If}$ | 0.52 | 6, 8 |
| Inhibitory baseline weight (co- & counter-tuned) | $w_{Ico}$ | 2.43 | 8, S2 |
| Inhibitory baseline weight (co- & counter-tuned) | $w_{Icounter}$ | 0.53 | 8, S2 |
| Noise parameter for inhibitory weights | $\epsilon_I^*$ | 0.01 | 4, 6, 8, S2 |
| Inhibitory baseline weight (Hebbian & scaling) | $w_{IF}$ | 0.8 | 5 |
| Noise parameter for inhibitory weights (Hebbian & scaling) | $\epsilon_I^*$ | 0.3 | 5 |
| Inhibitory baseline weight (Hebbian & anti-Hebbian) | $w_{IF}$ | 0.6 | 7 |
| Noise parameter for inhibitory weights (Hebbian & anti-Hebbian) | $\epsilon_I^*$ | 0.01 | 7 |
| Inhibitory baseline weight | $w_{IF}$ | Varying | S1, S3 |
| Noise parameter for inhibitory weights | $\epsilon_I^*$ | 0.01 | S1, S3 |

### Plasticity models

In this work we used three different inhibitory synaptic plasticity (ISP) rules. We termed them *Hebbian*, *scaling*, and *anti-Hebbian*. Both Hebbian and anti-Hebbian plasticity rules are triggered by pre- and postsynaptic spikes, and depend on a low-pass filter of these spike trains. The presynaptic trace (low-pass filter) is given by

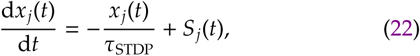

where *x*_*j*_(*t*) is the value of the trace of the spike train of presynaptic afferent *j* at time *t*; τ_STDP_ is the time constant of the trace, and *S*_*j*_(*t*) is a sum of Dirac delta functions (Eq. 4) representing the spike train of afferent *j*. The same is considered for the postsynaptic neuron,

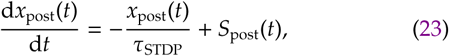

where *x*_post_(*t*) is the postsynaptic trace, and *S*_post_(*t*) is the spike train of the postsynaptic neuron (Eq. 5). Note that we used the same time constant for both pre- and postsynaptic traces.

#### Hebbian inhibitory plasticity

Precise balance of excitatory and inhibitory inputs was learned by a Hebbian inhibitory plasticity rule^6^. The weight of the jth inhibitory synapse changes according to

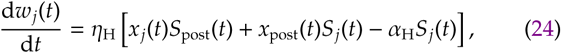

where *η*_H_ is the learning rate, and *α*_H_ is a parameter that defines the postsynaptic firing-rate. The first two terms on the right-hand side of Eq. 24 are Hebbian terms that increase the weights when both pre- and postsynaptic activities are correlated. The last term on the right-hand side of Eq. 24 is a penalty term for inhibitory spikes alone, which creates a homeostatic set-point for the postsynaptic firing-rate given by

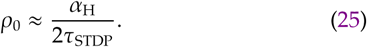

Later we describe how to arrive at this approximation.

#### Inhibitory synaptic scaling for flat tuning

One of the synaptic weight profiles we used for inhibitory synapses was flat, i.e., every synapse group had the same strength. To learn the flat profile from random initial weights we implemented a scaling plasticity rule, partially based on experimental work that observed synaptic scaling on inhibitory synapses^48,49^. Weights are increased if the postsynaptic firing-rates are too high, and decreased otherwise,

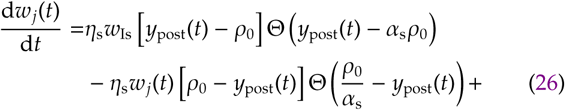

where *η*_s_ is a learning rate, ρ_0_ is a firing-rate reference value, chosen to be the same as the one for Hebbian plasticity rule,Θ(⋅) is the Heaviside function and *α*_s_ is a term that sets the firingrate range for which synapses do not change. Postsynaptic neuron’s firing-rate is computed with a slow averaging of the postsynaptic activity through

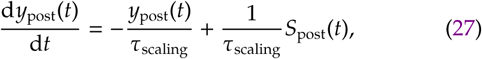

where τ_scaling_ is the time constant for the postsynaptic activity and *S*_post_(*t*) is the postsynaptic spike train (Eq. 5). Note that the last term on the right-hand side of the equation above is divided by τ_scaling_ so that *y*_post_(*t*) is in units of rate. Synaptic depression is weight dependent while synaptic potentiation is not, which ensures that all synaptic weights tend to the same value. When the postsynaptic neuron is firing below a threshold ρ_0_/*α*_s_, all inhibitory synapses in the flat group have their weights decreased proportionally to the difference between the target firing-rate and the average firing-rate, but also proportional to the current weight value. This way, strong synapses undergo stronger decrease than weak ones. Conversely, when the postsynaptic neuron is firing above a threshold *α*_s_ρ_0_, the same synapses increase in value by the same amount. Intuitively, these mechanism ensures that all synapses converge to the same value for a long run.

#### Anti-Hebbian inhibitory plasticity

The third inhibitory plasticity rule we used is an anti-Hebbian rule based on experimental data^21–23^ and theoretical work on recurrent networks^24^. Synaptic weights change according to

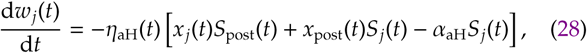

where *η*_aH_(*t*) is a variable learning rate and *α*_aH_ is a parameter to counterbalance the anti-Hebbian term. The resulting rule dictates that coincident events decrease inhibitory synapses, while non-coincident ones increase synaptic weights. Due to the unstable nature of this plasticity rule (see details below), we implemented a time-varying learning rate which evolves according to

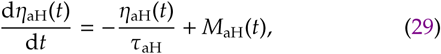

where τ_aH_ is the decay time constant for the learning rate, and *M*_aH_(*t*) is an external signal to transiently activate plasticity. We speculate that such signal could come from modulatory neurons such as dopaminergic or cholinergic and assumed that the external signal peaks at a time *t*_0_ (beginning of the simulation), so that

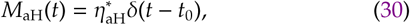

where *η*^∗^_aH_ is the maximum learning rate before decaying to zero, and *t*_0_ is the time when plasticity at these synapses are initiated. Parameters used for plasticity models are detailed in Table IV.

**TABLE IV.** Simulation parameters for plasticity rules.

| Parameter | Symbol | Value | Figs. |
| --- | --- | --- | --- |
| STDP time constant | $\tau_{STDP}$ | 20 ms | 5, 7, S1 |
| Hebbian learning rate | $\eta_H$ | $10^{-3}$ | 5, 7, S1 |
| Hebbian decay term | $\alpha_H$ | 0.2 | 5, 7, S1 |
| Firing-rate set-point | $\rho_0$ | 5 Hz | 5, 7, S1 |
| Scaling time constant | $\tau_{scaling}$ | 1000 ms | 5 |
| Scaling learning rate | $\eta_s$ | $5 \times 10^{-7}$ | 5 |
| Scaling learning rate weight | $w_{Is}$ | 0.6 | 5 |
| Scaling threshold parameter | $\alpha_s$ | 2 | 5 |
| Anti-Hebbian initial learning rate | $\eta_{aH}^*$ | $10^{-3}$ | 7 |
| Anti-Hebbian learning rate time constant | $\tau_{aH}$ | 200 s | 7 |
| Anti-Hebbian increase term | $\alpha_{aH}$ | 0.6 | 7 |
| Anti-Hebbian peak time | $t_0$ | 0 ms | 7 |
| Simulation time | – | 30 mins | 5, 7, S1 |

### Mean-field analysis of the plasticity rules

We were interested in plasticity rules with stable dynamics. For a better intuition on fixed-point dynamics and stability we consider here a simplified dynamics of a mean-field model for both the Hebbian^6^ and the anti-Hebbian models. We define the post-synaptic firing-rate as **ν**_post_(*t*) and the presynaptic firing-rates as *ν*_*j*_(*t*). The traces of both presynaptic afferent and postsynaptic neuron thus have an average of τ_STDP_*ν*_*j*_(*t*) and τ_STDP_*ν*_post_(*t*), respectively^68^. Neglecting any correlation between pre- and postsynaptic spikes, the average weight change for Hebbian synapses is given by

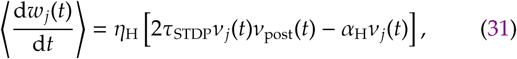

where 〈⋅〉s represents average over time. Intuitively, the post-synaptic firing-rate, *ν*_post_(*t*), changes negatively with changes in inhibitory weights – increased inhibition generates fewer post-synaptic spikes and vice-versa for decreased inhibition. This means that average firing rates are inversely linked to average inhibitory weights, i.e.,

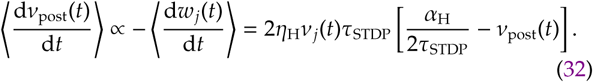

The steady state is computed by considering the vanishing point of the equation above (we assume that the presynaptic activity is nonzero), thus

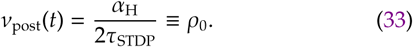

This means that the postsynaptic activity *ν*_post_(*t*) increases (via decreases in inhibitory efficacy) when below ρ_0_ and decreases when above ρ_0_, creating a stable fixed-point for the postsynaptic firing-rate.

The opposite is true for the anti-Hebbian plasticity rule. Changes in postsynaptic firing-rate (with the same assumption as for the Hebbian plasticity rule) follow

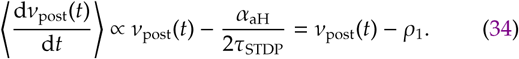

Because postsynaptic activity increases when it is above threshold ρ_1_ and decreases when it is below, this rule is unstable. The postsynaptic firing-rate eventually explodes or vanishes. We chose the simplest way to overcome these problems by setting a time-varying learning-rate. Other intricate mechanisms could be implemented, but this is not the scope of our work.

### Convergence of weights following the scaling plasticity rule

Our scaling plasticity rule has two different mechanisms, one for long-term depression (LTD) and one of long-term potentiation (LTP): LTD is multiplicative and LTP is additive (Eq. 26). The combined effect ensures that all incoming weights collapse to the same value (synaptic changes do not depend on presynaptic activity either). Here we present a mathematical intution to explain how synaptic weights can converge to the same value. First, we rewrite the scaling plasticity rule into two simplified terms. We consider constant postsynaptic firing-rate during LTD, 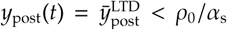. Consequently, the LTD part is described by

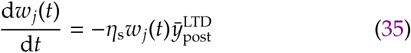

with solution

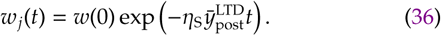

Doing the same for LTP 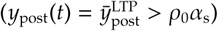 we arrive in

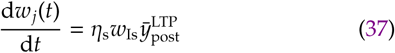

with solution

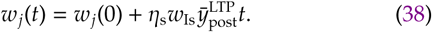

Defining 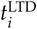 as the ith interval in which LTD occurred and 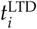 as the ith interval in which the synapse underwent LTP we can combine Eq. 36 and Eq. 38 (after a few lines of math) to rewrite the synaptic strength, *w*_*j*_(*t*), at time *t* as

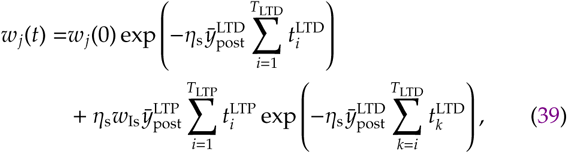

where *T*_LTD_ and *T*_LTP_ are the number of intervals with LTD and LTP, respectively. Because synaptic scaling is a continuous process, we can assume that LTP is always followed by LTD, and vice-versa, and thus *T*_LTD_ = *T*_LTP_ ± 1. Taking into consideration that the postsynaptic neuron’s firing-rate fluctuates around the target firing-rate, ρ_0_, LTD and LTP occur with similar rates,

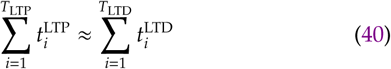

The first term on the right-hand side of Eq. 39 vanishes for long times, and the second term dominates with the late terms (*i* » 1, or, from *T*_LTP_ − κ to *T*_LTP_ for small κ),

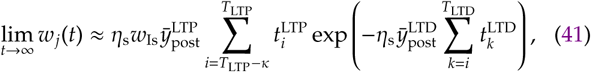

which is finite given that the postsynaptic neuron’s firing-rate fluctuates around the target firing-rate, ρ_0_. The final (stable) weight depends on the time the system stays in LTD and LTP after stabilisation, but not on the initial weight *w*_*j*_(0).

### Correlation

We quantified the response of the postsynaptic neuron to natural inputs with the Pearson correlation between postsynaptic firing-rate and input firing-rate fluctuations, per signal group. We computed the firing rate of a signal groups as the low-pass filter of the spike trains of its excitatory afferents,

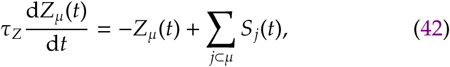

where *Z*_*µ*_(*t*) is the firing rate of the signal group *µ* at time *t*, filtered with a time constant τ_*Z*_. The postsynaptic activity is also computed through a low-pass filter of its spike train,

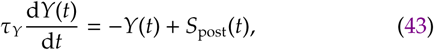

where *Y*(*t*) is the activity of the postsynaptic neuron at time *t* filtered with a time constant τ_*Y*_. The correlation is then computed as

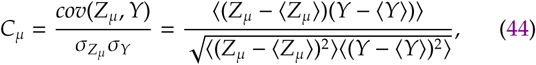

where *cov*(*z*, *y*) is the covariance between variables *z* and *y*, σ_*z*_ is the standard deviation of variable *z*, and 〈⋅〉 represents time average.

Subsequently we computed a *performance index* ∆*C* as the difference between the correlation measure for preferred (*µ* = 9) and non-preferred (*µ* = 1) input signals,

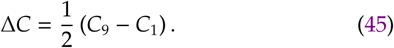

Maximum positive performance index, ∆*C* = 1, means that the preferred signal group has maximum correlation (*C*_9_ = 1) while the non-preferred signal group has maximum anti-correlation (*C*_1_ = − 1), indicating that the postsynaptic neuron is responding solely to the preferred signal group. Consequently, ∆*C* = −1, indicates that the postsynaptic neuron is responding solely to the non-preferred signal group. A flat response is indicated by ΔC = 0. Note that maximum ΔC (either positive and negative) is only achievable if there is no overlap between activation of preferred and non-preferred input signals, which is never the case here. We define as *best performance* when ∆*C* = 0 for all inhibitory inputs active (control), ∆*C* = 1 (or ∆*C* > 0) for one inhibitory population inactive, and ∆*C* = −1 (or ∆*C* < 0) when the other inhibitory population is inactive. Parameters used for computing correlations are detailed in Table V.

**TABLE V.** Simulation parameters for correlation measure.

| Parameter | Symbol | Value | Figs. |
| --- | --- | --- | --- |
| Presynaptic time constant | $\tau_Z$ | 10 ms | 4, 6, 8, S2, S3 |
| Postsynaptic time constant | $\tau_Y$ | 250 ms | 4, 6, 8, S2, S3 |
| Simulation time | – | 30 mins | 4, 6, 8, S2, S3 |

### Implementation

Models were simulated with a time-step ∆*t*, with either analytical or *semi-analytical* solution of the corresponding differential equation. All codes were written in Fortran, compiled with Intel Fortran Compiler 19.0, running on an Intel-based Linux computer (Debian 9; i9-9900X processor; 32 GB memory). Bellow we describe how each equation was implemented, with the parameter values in tables in the end of this section.

When not in the refractory period (see below), the leaky integrate-and-fire neuron is updated as

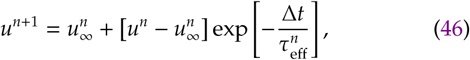

where *n* is the iteration index, 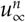 and 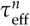 are auxiliary variables described by

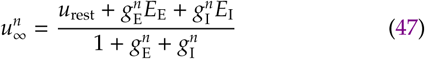

and

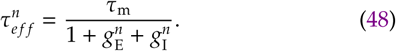

This is the analytical solution when considering that all variables apart from *u*(*t*) are constant during a time-step, which we refer to as *semi-analytical*.

When the membrane potential crosses a threshold from below, the membrane potential is reset (because of a spike being triggered), and kept at the reset potential for the duration of the refractory period,

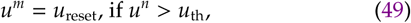

where

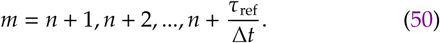

Synaptic conductances are implemented as

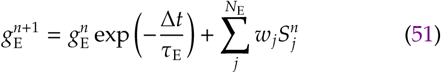

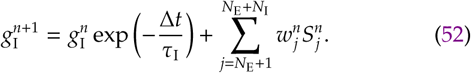

Note that here 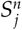 is equal to one when afferent *j* spiked at time-step *n* and zero otherwise.

For natural inputs, we updated the auxiliary variable *y*_*µ*_(*t*) every Δ*T*,
an action potential is

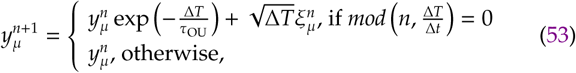

where *mod*(⋅, ⋅) is the modulo operation and 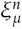 is a random number drawn from a gaussian distribution with zero mean and unitary standard deviation. Presynaptic spikes are generated as point processes, so that at each time-step the probability of a presynaptic afferent to spike is

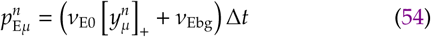

and 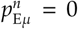 during the τ_Eref_/∆*t* iterations after a spike. The same is valid for an inhibitory afferent; the probability of firing an action potential is

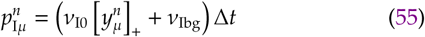

and 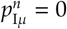 during the τ_Iref_/∆*t* iterations after a spike.

For pulse inputs, presynaptic afferents were set to fire at background firing-rate and had an elevated firing-rate during a 100-ms period, which was varied in 5 Hz steps. For the activated pattern

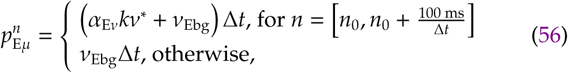

where *α*_E*ν*_ adjusts the excitatory firing-rate, *k* is an integer for varying the pulse intensity, and *n*_0_ is the iteration in which the pulse starts. The same implementation was used for inhibitory afferents (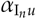 being the parameter to adjust the inhibitory firing-rate),

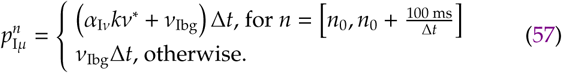

An afferent in an inactive pattern fires action potentials with background frequency (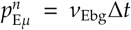 and 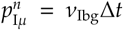), and there is no spike elicit in the refractory period (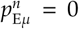 and 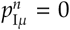 during the τ_Eref_/Δ*t* and τ_Iref_/Δ*t* iterations after a spike, respectively).

Plasticity was implemented with spike triggered events. For the Hebbian and anti-Hebbian plasticity rules, auxiliary variables changed as

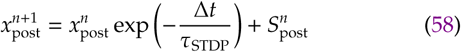

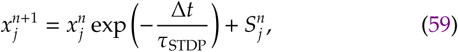

where 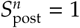 if if the postsynaptic neurons generated an action potential at iteration *n* and zero otherwise. Hebbian weights changed according to

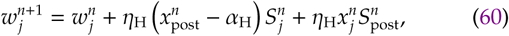

and anti-Hebbian to

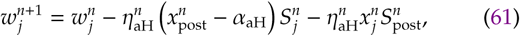

with the learning rate varying as

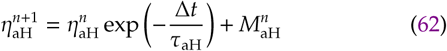

with 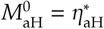, and 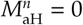 for *n* > 0.

Scaling was implemented with a different trace,

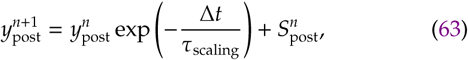

with weight update following

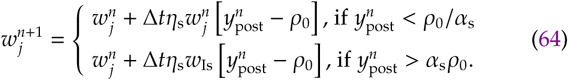

Correlation-related variables were updated as

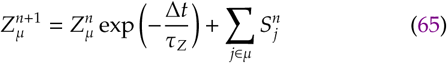

and

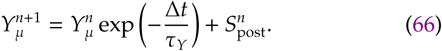

## Supplementary figures

**FIG. S1.**
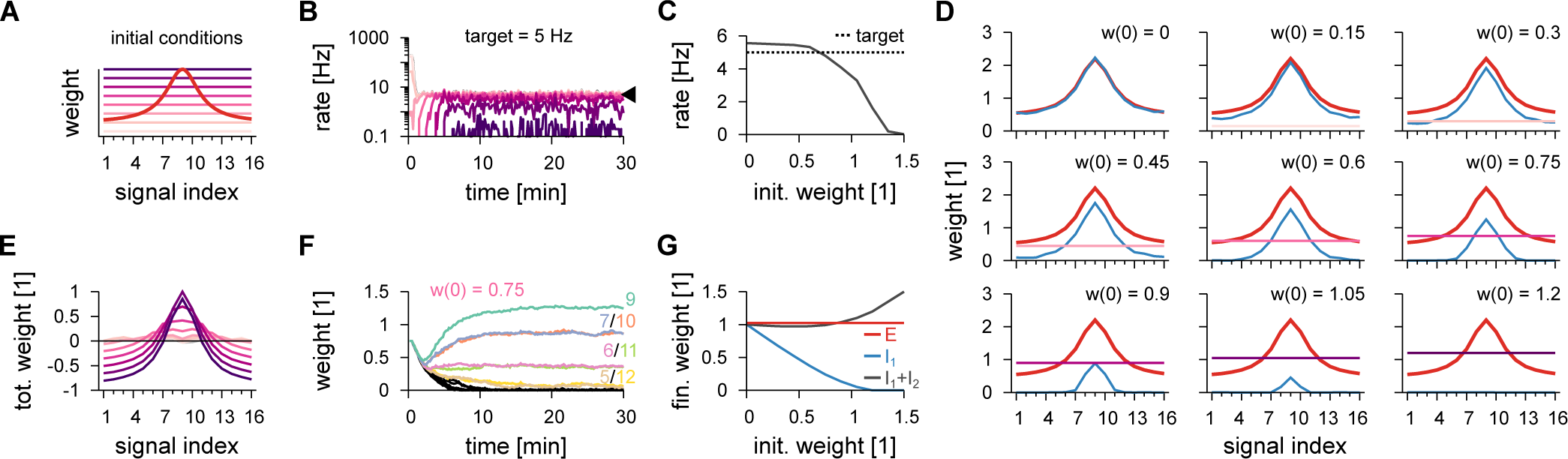
Inhibitory plasticity acting on one inhibitory population compensates global inhibition from a second inhibitory population. **A**, Schematic of the synaptic weight profile for excitatory synapses (red) and different initial conditions for inhibitory synapses (pink to purple colour-code). Inhibitory population 1 has its inhibitory synapses changing according to a plasticity mechanism while population 2 remains fixed. **B**, Time-course of the postsynaptic firing-rate for different initial conditions (colours as in A). Inhibitory plasticity on population 1 is set to achieve a balanced state with target of 5 Hz (arrowhead). **C**, Stabilised postsynaptic firing-rate as a function of the initial inhibitory synaptic weight. **D**, Individual synaptic weight profiles for excitatory (red), inhibitory population 1 (blue, after synaptic stabilisation), and inhibitory population 2 (colour coded as A). **E**, Total synaptic weight per signal (excitatory minus inhibitory) for different initial conditions after stabilisation of synapses from population 1. **F**, Example of synaptic dynamics of inhibitory population 1 for a given initial condition. Colours represent different signal groups. **G**, Final weights as a function of initial inhibitory weights. Plotted are excitatory (red), plastic inhibitory (blue) and sum of total inhibitory synapses (grey).

**FIG. S2.**
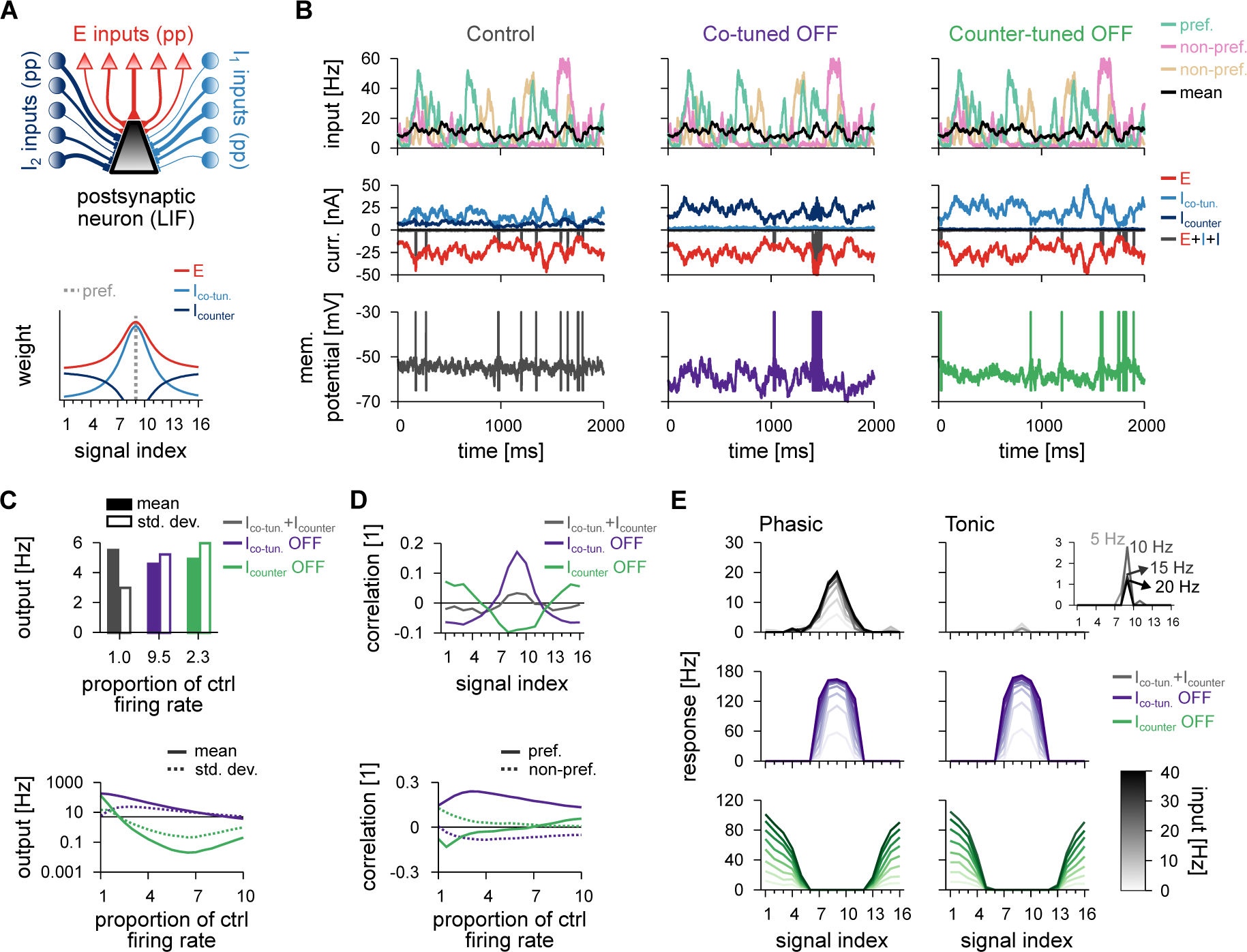
Postsynaptic response for the model with co- and counter-tuned inhibitory populations. **A**, Schematic of the circuit with two inhibitory populations (top); *I*_1_ corresponds to co-tuned and *I*_2_ to counter-tuned population. Pre-synaptic spikes were generated as point-processes (pp) and fed into an LIF. Synaptic weight profile (bottom). Average weight (y-axis) for different input signals (x-axis); preferred signal is pathway no. 9 (grey dashed line). **B**, Average firing-rate of the preferred and two non-preferred inputs and mean of all inputs (top row), total excitatory current and inhibitory currents of both populations (middle row), and membrane potentials (bottom row), for control (left), co-tuned (middle) and counter-tuned (right) population inactive. **C**, Average and standard deviation of the postsynaptic firing-rate due to natural input for the three cases (top), and as a function of the inhibitory firing-rate (bottom). **D**, Pearson correlation between postsynaptic firing-rate and excitatory input firing-rates for different input signals for the three conditions in B. Correlation between preferred (continuous line) or non-preferred (dashed line) with the output activity as a function of the inhibitory firing-rate of each inhibitory population (bottom). **E**, Response to a pulse input in the phasic (left; first 50 ms), and tonic (right; last 50 ms) periods. Firing rate computed as the average number of spikes (for 100 trials) normalised by the bin size (50 ms). Each line corresponds to a different input strength; from light (low amplitude pulse) to dark (high amplitude pulse) colours. Insets show tonic response for control firing-rates.

**FIG. S3.**
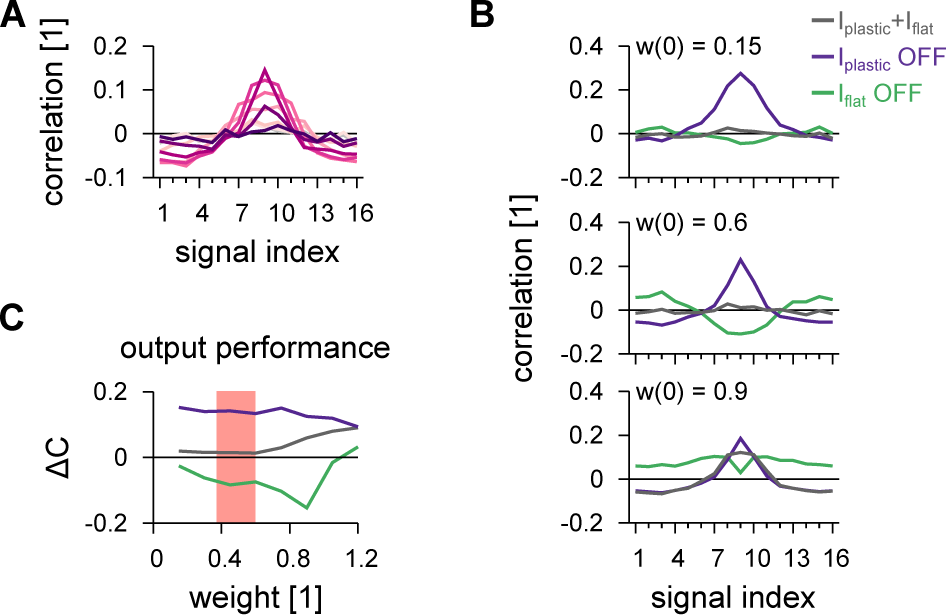
Postsynaptic response after stabilisation of synapses from one population. **A**, Pearson correlation between postsynaptic firing-rate and excitatory input firing-rates with both inhibitory populations active. Colour code as in Fig. S1A. **B**, Pearson correlation between postsynaptic firing-rate and excitatory input firing-rates for different signals with both inhibitory populations active, population 1 inactive, and population 2 inactive. Three examples are shown. **C**, Performance index as the difference between preferred signal and non-preferred signal as in Fig. 8A, but for the whole range of different initial conditions (inhibitory synaptic weights). Pink shaded area corresponds to best performance, which maximises the difference between preferred and non-preferred correlation responses oppositely for inactivation of different populations, and minimising difference between preferred and non-preferred correlation when both inhibitory populations are active.

